# Simultaneous pure T_2_ and varying T_2_′-weighted BOLD fMRI using Echo Planar Time-resolved Imaging for mapping cortical-depth dependent responses

**DOI:** 10.1101/2021.05.22.445292

**Authors:** Fuyixue Wang, Zijing Dong, Lawrence L. Wald, Jonathan R. Polimeni, Kawin Setsompop

**Affiliations:** Athinoula A. Martinos Center for Biomedical Imaging, Massachusetts General Hospital, Charlestown, Massachusetts, USA; Harvard-MIT Health Sciences and Technology, MIT, Cambridge, Massachusetts, USA; Electrical Engineering and Computer Science, MIT, Cambridge, Massachusetts, USA; Department of Radiology, Harvard Medical School, Boston, Massachusetts, USA; Department of Radiology, Stanford University, Stanford, USA; Department of Electrical Engineering, Stanford University, Stanford, USA

**Keywords:** high field, multi-echo, spin-echo, T_2_ BOLD, microvascular specificity, laminar fMRI

## Abstract

Spin-echo (SE) BOLD fMRI has high microvascular specificity, and thus provides a more reliable means to localize neural activity compared to conventional gradient-echo BOLD fMRI. However, the most common SE BOLD acquisition method, SE-EPI, is known to suffer from T_2_′ contrast contamination with undesirable draining vein bias. To address this, in this study, we extended a recently developed distortion/blurring-free multi-shot EPI technique, Echo-Planar Time-resolved Imaging (EPTI), to cortical-depth dependent SE-fMRI at 7T to test whether it could provide purer SE BOLD contrast with minimal T_2_′ contamination for improved neuronal specificity. From the same acquisition, the time-resolved feature of EPTI also provides a series of asymmetric SE (ASE) images with varying T_2_′ weightings, and enables extraction of data equivalent to conventional SE EPI with different echo train lengths (ETLs). This allows us to systematically examine how T_2_′-contribution affects different SE acquisition strategies using a single dataset. A low-rank spatiotemporal subspace reconstruction was implemented for the SE-EPTI acquisition, which incorporates corrections for both shot-to-shot phase variations and dynamic *B*_0_ drifts. SE-EPTI was used in a visual task fMRI experiment to demonstrate that i) the pure SE image provided by EPTI results in the highest microvascular specificity; ii) the ASE EPTI series, with a graded introduction of T_2_′ weightings at time points farther away from the pure SE, show a gradual sensitivity increase along with increasing draining vein bias; iii) the longer ETL seen in conventional SE EPI acquisitions will induce more draining vein bias. Consistent results were observed across multiple subjects, demonstrating the robustness of the proposed technique for SE-BOLD fMRI with high specificity.

## 1. Introduction

Gradient-echo (GE) blood oxygenation level-dependent (BOLD) with T_2_^*^ weighting is one of the most commonly used fMRI contrasts due to its high sensitivity and acquisition efficiency (Bandettini et al., 1992; Kwong et al., 1992; Ogawa et al., 1990). However, the GE BOLD signal contains a mixture of contributions from both large and small blood vessels. The macrovascular signals, such as those from large draining veins, can be far from the site of neuronal activity (Havlicek and Uludağ, 2020; Heinzle et al., 2016; Markuerkiaga et al., 2016). Therefore, the inclusion of macrovascular signal in GE BOLD fMRI significantly limits its effective resolution to detect brain activity, even at high spatial resolution at ultra-high-field.

Conversely, the microvascular signal can provide more specific and precise localization of the neuronal activity (Dumoulin et al., 2018; Norris and Polimeni, 2019), therefore, a number of alternative fMRI contrasts have been investigated that have the promise of achieving high neuronal specificity through being sensitive to signal changes from the microvasculature while being insensitive to changes from the macrovasculature (De Martino et al., 2018; Huber et al., 2019; Koopmans and Yacoub, 2019). Among those, spin-echo (SE) or T_2_ BOLD fMRI has shown great potential (Bandettini and Wong, 1995; Boxerman et al., 1995; Ogawa et al., 1993) and has been demonstrated to provide superior specificity compared to GE BOLD (Huber et al., 2017a). However, it is difficult to efficiently obtain pure T_2_-weighting without T_2_′ contamination (Norris, 2012), which introduces unwanted sensitivity to the microvasculature and thus compromises its achievable neuronal specificity.

This difficulty originates from the technical challenges in conventional MRI acquisition. For example, the most commonly-used T_2_ BOLD acquisition method, spin-echo EPI, uses a long echo-train-length (ETL) that samples both T_2_- and T_2_′-weighted signals to generate an image, and therefore suffers from T_2_′ contamination with an undesirable sensitivity to large blood vessels (Bandettini et al., 1994; Birn and Bandettini, 2002; Duyn, 2004; Goense and Logothetis, 2006; Koopmans and Yacoub, 2019; Norris, 2012; Yacoub et al., 2003). In-plane acceleration with parallel imaging (Griswold et al., 2002; Pruessmann et al., 1999a; Sodickson and Manning, 1997) or multi-shot EPI (ms-EPI) (Butts et al., 1996; Chen et al., 2013; Dong et al., 2018; Jeong et al., 2013; Mani et al., 2017) can be used to address this issue by reducing the effective echo spacing and thus the ETL. However, their abilities to reduce T_2_′ contribution come at a cost of higher noise amplification or image artifacts due to shot-to-shot phase variations, and residual T_2_′ contamination as well as distortion and blurring still remains. Other alternative sequences have been proposed to provide T_2_ BOLD contrast (Barth et al., 2010; Bowen et al., 2005; Chamberlain et al., 2007; Constable et al., 1994; Denolin and Metens, 2003; Goerke et al., 2011; Miller et al., 2003; Polimeni et al., 2017; Poser and Norris, 2007; Scheffler et al., 2001). Among those, 3D-GRASE uses multiple refocusing pulses with short echo trains in between, which significantly reduces the T_2_′ weightings and offers higher specificity than conventional SE-EPI (Beckett et al., 2020; De Martino et al., 2013; Feinberg et al., 2008; Kemper et al., 2015; Olman et al., 2012; Park et al., 2021). Recent works have demonstrated its specificity for high-resolution SE BOLD fMRI at 7T (Beckett et al., 2020; Kemper et al., 2015), but challenges including high SAR, relatively small achievable coverage and a mixture of T_1_ weightings from stimulated echoes (Goerke et al., 2007) remain to be addressed or interpreted. In addition to SE BOLD fMRI, non-BOLD contrasts such as cerebral blood volume (CBV) has shown promising results as well, such as using the vascular space occupancy (VASO) methods (Chai et al., 2020; Huber et al., 2020; Huber et al., 2014; Huber et al., 2019; Jin and Kim, 2006; Lu et al., 2003; Lu et al., 2004).

In this study, to address the T_2_′-contamination of SE-EPI and obtain higher neuronal specificity with reduced draining vein effects, we extended a recently developed technique, Echo-Planar Time-resolved Imaging (EPTI) (Wang et al., 2019; Wang et al., 2020; Wang et al., 2021b), to high-resolution SE-fMRI at ultra-high-field for mapping laminar fMRI responses. EPTI is a novel multi-shot EPI approach that has been previously developed for efficient multi-contrast and quantitative mapping. It employs a novel spatiotemporal encoding strategy in the frequency-echo (*k*-*t*) domain, and is therefore able to resolve a series of multi-contrast images across the readout with a small TE increment as short as an echo spacing (∼1 ms). The images are also distortion- and blurring-free, providing accurate anatomical information for high-resolution imaging. In addition, the continuous signal readout scheme and the use of spatiotemporal correlation to recover the *k-t* undersampled data in EPTI result in high acquisition efficiency, allowing us to acquire multi-echo images at submillimeter isotropic resolution within a few shots.

Here, we show that the time-resolved feature of EPTI can not only provide pure T_2_ contrast images to increase microvascular specificity, but can also simultaneously acquire T_2_′-weighted images to investigate the macrovascular contribution across the spin-echo readout. Specifically, using a SE-EPTI acquisition, we obtained a pure SE image with minimal T_2_′ contamination, and a series of asymmetric SE (ASE) images with varying T_2_′ weightings. We also developed a framework to extract conventional SE ms-EPI images with different ETLs (therefore with different levels of T_2_′-contamination) from the same dataset, but without any distortion. This ensures that all images concurrently acquired in a SE-EPTI acquisition, including the pure SE, ASEs and the extracted conventional SE-EPIs are perfectly matched and aligned. A subspace reconstruction (Dong et al., 2020; Guo et al., 2021; He et al., 2016; Lam and Liang, 2014; Liang, 2007; Meng et al., 2021; Tamir et al., 2017) was further implemented for SE-EPTI datasets, which incorporates corrections for both dynamic *B*_0_ drifts and shot-to-shot phase variations caused by respiration or other physiological factors. A 3-shot SE-EPTI protocol was developed to acquire a thick slab that sufficiently covers the visual cortex at submillimeter resolution (0.9 mm isotropic). Through cortical depth analyses, we demonstrated that the pure SE BOLD data provided by EPTI results in the highest microvascular specificity as expected, and the ASE EPTI image series, with a graded introduction of T_2_′ weightings at time points farther away from the pure SE, show gradually increased sensitivity, but larger and larger draining vein bias. Using the same dataset, we also experimentally confirmed that a longer ETL in the conventional SE EPI acquisition will induce more draining vessel bias. We present consistent results across multiple subjects, demonstrating the robustness of the proposed technique and framework.

## 2. Material and methods

### 2.1. EPTI for pure contrast multi-echo imaging

Conventional EPI acquires one phase encoding position per readout line through a fast continuous bipolar readout, and forms a single image by combing all of these signals acquired at different time points. Although its image contrast is mainly determined by the “effective” echo time (TE), the time when the central PE line is acquired, signal decay occurs across the sampling window and contrast weightings from other time points also contribute to the final image. For SE-EPI acquisition, the signal magnitude *S*(*t*) at time *t* after excitation in a mono-exponential model is:

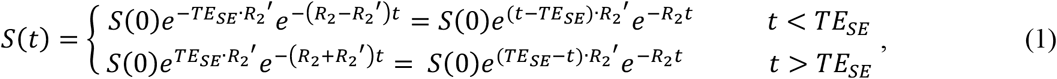

where *S*(0) represents the initial signal intensity at *t* = 0, *TE*_*SE*_ is the echo time of the spin-echo, *R*_2_ is the T_2_ relaxation rate, 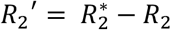, representing the difference between T_2_^*^ and T_2_ relaxation rates and reflecting the susceptibility*-*induced recoverable intra-voxel dephasing, which is sensitive to extravascular BOLD surrounding all vessels including large veins. As shown in the signal model, pure T_2_-weighted signal can be obtained at *TE*_*SE*_, while all the other time points will be affected by *R*_2_′*-*weighting with an extent determined by the time interval between their acquisition time and the SE refocusing time. Signals acquired at those time points with *R*_2_′-weighting will therefore induce contrast contamination in SE-EPI when they are combined to form an EPI image, leading to increased sensitivity to large vessels and compromised neuronal specificity.

The main concept of SE-EPTI is to recover a series of images across the readout at all those time points by recovering fully-sampled data across the *k*-*t* space (Dong et al., 2021a; Dong et al., 2020; Dong et al., 2019; Wang et al., 2019; Wang et al., 2021a). Figure 1a shows the sequence diagram of SE-EPTI that is used to achieve this goal, where an EPI-like continuous readout is used in a SE acquisition with different G_y_ gradient blips applied to sample a *k*_y_-segment in *k*-*t* space using a spatiotemporal CAIPI encoding pattern (Fig. 1b). This sampling pattern not only ensures that the neighboring *k*_y_ points are sampled within a short time interval with high temporal correlations, but also that they are interleaved and complementary along the *k*_y_ direction in an optimized pattern. It allows efficient use of temporal correlation and coil sensitivity to achieve high undersampling, therefore only a few EPTI shots are needed to cover the desired *k*-*t* space for imaging. After the full *k*-*t* space is recovered in the reconstruction, multi-contrast images with pure contrast at all time points can be simply obtained by an inverse Fourier transform, including a pure T_2_ SE image and a series of ASE images with varying T_2_′ weighting, spaced at a TE increment of an echo-spacing (∼1 ms) as shown in Fig. 1b.These images are free from any distortion and blurring artifacts, which are common in conventional EPI due to *B*_0_-inhomogeneity-induced phase accumulation and signal decay across the readout. This is because each EPTI image is recovered using signals with exactly the same phase and magnitude.

**Figure 1.**
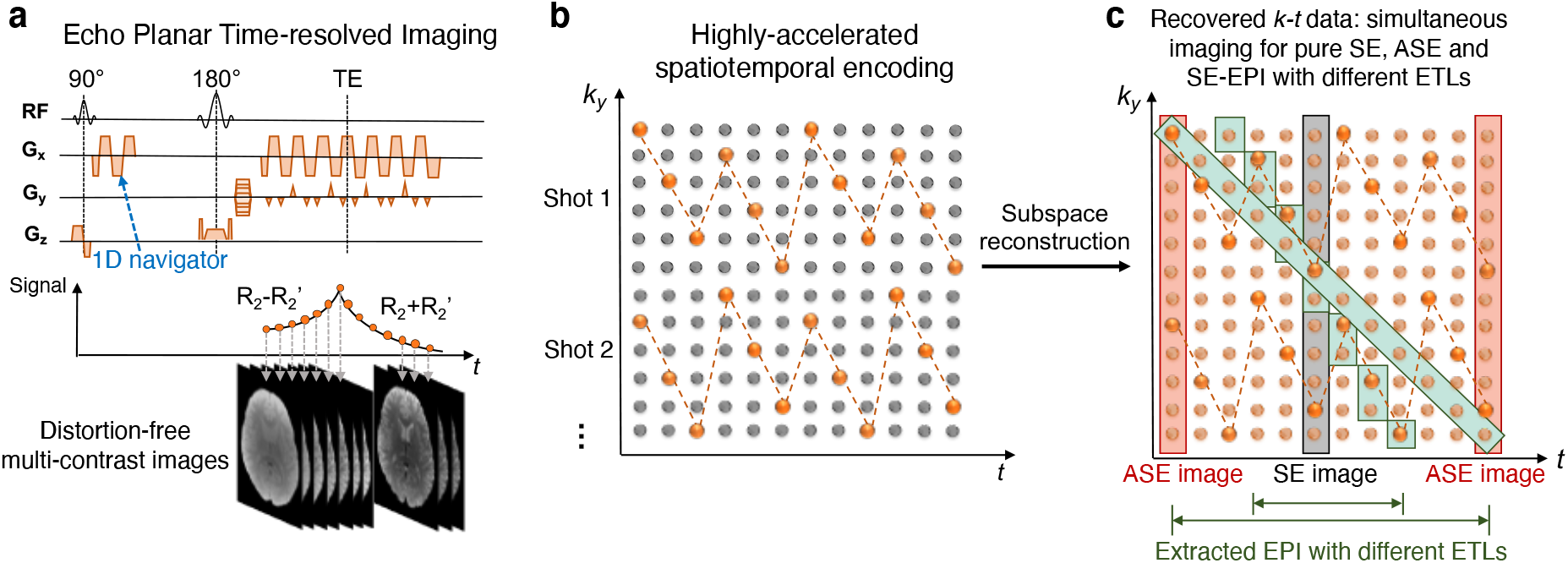
Spatiotemporal CAIPI encoding of EPTI and generation of multi-contrast images. The recovered *k-t* data after reconstruction can provide SE image with pure T_2_ weighting (gray), asymmetric SE (ASE) images with both T_2_ and T_2_′ weighting (orange), and extracted SE-EPI with different ETLs (green) to investigate the effect of T_2_′ contamination.

### 2.2. EPTI-extracted conventional EPI with different echo train lengths

In conventional SE-EPI acquisition, the reduction of ETL can help reduce the level of *R*_2_′ contamination, which can be achieved through in-plane acceleration using parallel imaging and/or through multi-shot segmentation. However, a large reduction factor in ETL would lead to either large noise amplification and aliasing artifacts in parallel imaging and/or a long acquisition time in multi-shot acquisition. Therefore, it remains challenging to achieve short ETLs for SE-EPI especially at high spatial resolution.

To investigate the benefit of reducing T_2_′ contamination by using a shorter ETL as well as comparing the conventional SE-EPI with the pure SE images, data that are equivalent to a conventional EPI-like acquisition with different ETLs can be extracted from the reconstructed *k*-*t* data acquired by EPTI. Fig. 1c shows a simplified illustration of such extraction, where reconstructed *k*-space signals at different time points or TEs are extracted in a diagonal pattern in *k*-*t* space. To mimic an interleaved ms-EPI acquisition, multiple adjacent PE lines are extracted at each time point depending on the shot number (e.g., 4 PE lines for a 4-shot EPI), and the final ETL of the extracted SE ms-EPI is determined by the overall matrix size along PE direction as well as the shot number. For example, for a 4-shot EPI acquisition with a matrix size of 144, 4 PE lines are extracted at each time point, and the resultant ETL will be 36. Before the extraction, *B*_0_-induced phase is removed from the *k*-*t* data by removing the TE-dependent linear phase changes in the image domain, so that the EPTI-extracted SE-EPI images are also free from any distortion. This ensures that all the image contrasts obtained concurrently from a single EPTI dataset, including the pure SE, the ASE series and the extracted SE-EPI, are geometrically matched, allowing for reliable evaluation of the impact of T_2_′ contamination with different ETLs on the signal contributions.

### 2.3. Subspace image reconstruction

The reconstruction framework of EPTI used in this study is shown in Fig. 2. To reconstruct the image series from the undersampled *k-t* data, a low-rank subspace method (Dong et al., 2020; Guo et al., 2021; He et al., 2016; Lam and Liang, 2014; Liang, 2007; Meng et al., 2021; Tamir et al., 2017) is applied to resolve signal evolutions across different TEs using signal model priors (Fig. 2c). Here, the reconstruction is performed to estimate a small number of coefficient maps of pre-calculated temporal subspace bases that can accurately represent the signal evolution across echoes, rather than to estimate all of the echo images directly. Such a reconstruction approach can achieve improved reconstruction conditioning and good reconstruction accuracy by taking advantage of the high spatiotemporal correlation in the EPTI datasets. In this work, we tailor this subspace reconstruction specifically to the SE-EPTI acquisition and further incorporate corrections for phase variations due to *B*_0_ changes across different shots and dynamics, increasing the robustness and accuracy of the reconstruction for fMRI experiments.

**Figure 2.**
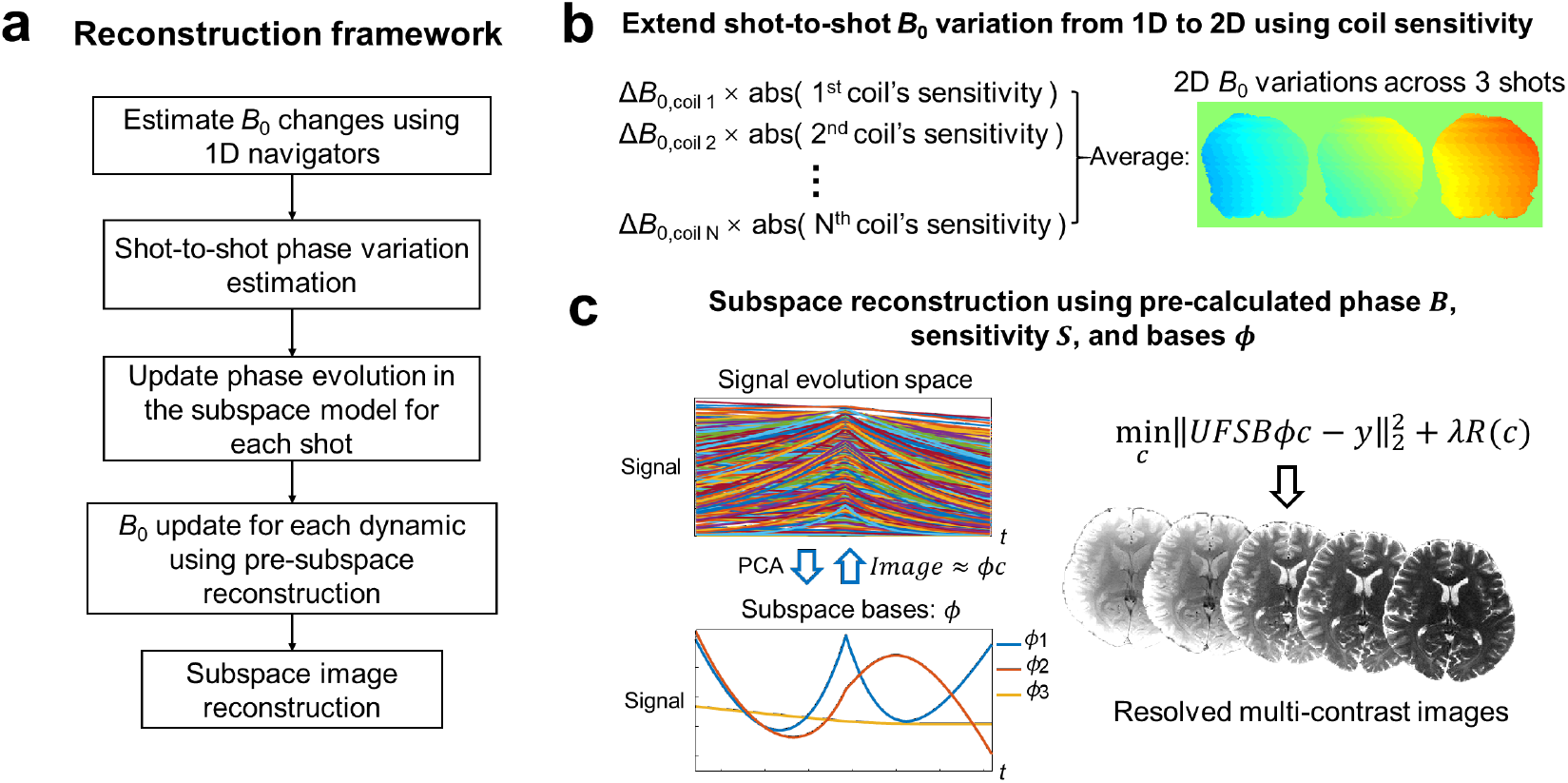
(a) EPTI reconstruction framework with dynamic *B*_0_ correction and shot-to-shot phase variation correction. (b) Illustration of the estimation method for shot-to-shot *B*_0_ variation using multi-channel 1D navigators. (c) Subspace reconstruction to resolve multi-contrast images by solving a small number of coefficient maps of the subspace bases.

In the reconstruction, the first step is to use principal component analysis (PCA) to generate a group of subspace bases *ϕ* from simulated signal evolutions across a range of possible T_2_ and T_2_^*^ values. The number of bases is selected to approximate all the simulated signal evolutions accurately with an error of <0.2% (3 bases in this work). Then, the coefficient map of the basis, *c*, is estimated by:

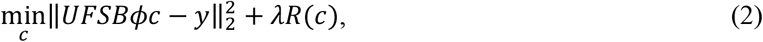

Here, *B* is the image phase evolutions across echoes caused by field inhomogeneity, *S* is the sensitivity map, *F* denotes the Fourier transform, *U* represents an undersampling mask, and *y is t*he acquired undersampled data. *R* is the locally low-rank (LLR) constraint applied on the coefficients to further improve the conditioning, and *λ* is the control parameter. The image phase evolution *B* and coil sensitivity *S* are estimated from a low-resolution *k-t* calibration scan acquired prior to the imaging scan. After estimating the coefficients, multi-contrast echo images can be recovered by performing a temporal expansion (*ϕc*) for all the TEs in each volume TR.

### 2.4. Shot-to-shot phase variation correction using navigator

The *B*_0_-inhomogeneity-induced phase accumulation could change temporally due to *B*_0_ drift and/or respiratory motion. Instead of inducing aliasing artifacts as in the conventional interleaved ms-EPI, such phase variations were shown to only cause minor local image smoothing on EPTI images due to the *k*_*y*_ block-segmented sampling pattern used (Wang et al., 2019). To mitigate these smoothing effects and improve the reconstruction accuracy, a method to estimate and correct for such shot-to-shot phase variations is incorporated into the reconstruction framework as shown in Fig. 2.

One dimensional (1D) *B*_0_ changes along *x* between all of the EPTI-shots and temporal dynamics are calculated first. Specifically, the phases of the 1^st^ and 3^rd^ echo of the standard 3-line EPI phase correction navigator (acquired after the excitation pulse but before the refocusing) are subtracted and scaled based on their TEs to obtain the *B*_0_ of every TR, and then the relative *B*_0_ changes to the first TR are calculated. The 1^st^ and 3^rd^ echo are used in this calculation to avoid odd-even echo phase difference in the bipolar readout. These 1D *B*_0_ changes are then extended to 2D by using the spatial information provided by the multi-channel coil sensitivities (Splitthoff and Zaitsev, 2009; Versluis et al., 2012; Wallace et al., 2020). Since the shot-to-shot *B*_0_ change varies smoothly in the spatial domain, the low-frequency spatial information provided by the multi-channel coils should be sufficient in capturing and recovering its spatial distribution along the PE direction. Specifically, the 2D *B*_0_ change, *ΔB*_2*D*_, can be approximated by a weighted combination of the 1D *B*_0_ changes from different coils using the magnitude of their coil sensitivities, similar to a previous approach (Versluis et al., 2012). The estimation process is as follows:

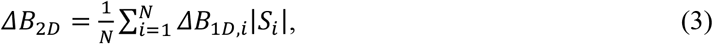

where *ΔB*_1*D,i*_ is the 1D *B*_0_ change estimated from the *i*-th coil, and |*S*_*i*_| is the magnitude of the *i*-th coil’s sensitivity map. Next, to correct for the shot-to-shot *B*_0_ changes in the reconstruction, the phase evolution term *B* in Eq. 2 is updated to incorporate the above estimated *B*_0_-induced phase into the forward model. A simulation experiment was performed to evaluate the effectiveness of the above shot-to-shot phase variation correction approach. The simulated data were generated using the imaging parameters and a set of quantitative maps (T_2_, T_2_^*^, *B*_0_ map) obtained from EPTI data in the in-vivo experiments, with additional smooth *B*_0_ changes (spatially 2^nd^ order) added to each shot as described below.

To account for higher-order *B*_0_ spatial changes, a pre-reconstruction process is also implemented to update the *B*_0_ maps across different dynamics. It uses pre-reconstructed 2D phase maps to help better estimate higher-frequency spatial variations of *B*_0_ changes, which can be used to adjust the forward model to improve the final reconstruction accuracy. In the *B*_0_ update pre-reconstruction, a set of complex subspace bases are first extracted from simulated signals with a range of *B*_0_ changes, which are then used in the pre-reconstruction model to estimate higher order phase evolution and *B*_0_. The estimated *B*_0_ changes are filtered by a hamming filter to remove any noise and potential artifacts, and then incorporated into the final image reconstruction by updating the phase term *B* (Eq. 2), similar to the approach described in (Dong et al., 2021a). More number of bases were used in the *B*_0_ update pre-reconstruction than in the image reconstruction (6 complex bases vs. 3 real bases) to provide additional degrees of freedom to model and estimate large *B*_0_ phase evolution.

### 2.5. Data acquisition

Written informed consent was obtained from each participant before the experiment in accordance with our institution’s Human Research Committee. All data were acquired on a Siemens Magnetom Terra 7T scanner (Siemens Healthineers, Erlangen, Germany), using a custom-built 64-channel receiver coil (Mareyam et al., 2020) with a single RF transmission channel.

SE-EPTI data were acquired on 3 healthy volunteers using the following acquisition parameters: FOV = 218 × 130 × 25.2 (RO × PE × slice, HF-LR-AP) mm^3^, matrix size = 240 × 144 × 28, 0.9-mm isotropic resolution, number of EPTI-shots (segmentation) = 3, number of echoes = 45, TE_range_ = 40–88 ms, TE of SE = 64 ms, echo spacing (TE-increment) = 1.09 ms, volume TR = 3 s × 3-shot = 9 s, 43 dynamics, acquisition time per run = 6 min 27 s, 14 runs were acquired for each subject. A standard block-design “checkerboard” visual stimulus contrast-reversing at 8 Hz was presented for the fMRI acquisitions. An initial 27-s fixation period was performed followed by four 36 s-54 s on-and-off blocks. To assist with fixation, a red dot with time-varying brightness was positioned at the center of the screen, and the subjects were asked to press a button as soon as they detected a change in its brightness. Before the EPTI data acquisition in each run, a fast *k-t* calibration scan was acquired in 54 s with a matrix size of 240 × 49 × 28 (RO × PE × slice) and 7 echoes. For each volunteer, a multi-echo magnetization-prepared rapid gradient echo (MEMPRAGE) (van der Kouwe et al., 2008) image was acquired at 0.75-mm isotropic resolution as an anatomical reference with a FOV of = 218 × 168 × 194 (AP-LR-HF) mm^3^.

Conventional single-shot SE-EPI and GE-EPI were also acquired for comparison on one of the healthy volunteers. The acquisition parameters for SE-EPI were: FOV = 218 × 130 × 25.2 (RO × PE × slice, HF-LR-AP) mm^3^, matrix size = 240 × 145 × 28, 0.9-mm isotropic resolution, TE = 64 ms, GRAPPA factor = 3, ETL = 52 ms, echo spacing = 1.09 ms, TR = 3 s. The GRAPPA factor of 3 was used here to allow the same TE to be achieved as the EPTI acquisition. The GE-EPI used the same FOV and resolution, and other acquisition parameters were: TE = 28 ms, GRAPPA factor = 4, ETL = 39 ms, echo spacing = 1.09 ms, TR = 3 s. Noted that a GRAPPA factor of 3 or 4 used here is already high, considering the small FOV along PE in our acquisition (i.e., ∼ 1.6x zoomed compared to standard axial scan with PE along AP). 129 dynamics were acquired per run for both GE- and SE-EPI in an acquisition time of 6 min 27 s. 4 runs were acquired for both GE- and SE-EPI. In order to estimate field maps and correct for geometric distortions, PE-reversed data were acquired for both GE- and SE-EPI with matched acquisition parameters prior to the fMRI acquisitions. The standard GRAPPA (Griswold et al., 2002) reconstruction was performed followed by complex coil combination (Pruessmann et al., 1999b). In addition, a turbo spin-echo (TSE) image was acquired with the same FOV and matrix size as a distortion-free reference.

### 2.6. Image post-processing

To align all the volumes from different dynamics and runs, registration was performed using AFNI (Cox, 1996). For EPTI, the motion parameters were estimated using the all-echo-averaged volumes with higher SNR, which were then applied to different multi-echo volumes as well as the extracted ms-EPI images. After registration, the multi-run data were averaged to a single dataset. For ss-EPI, distortion correction was performed using the *B*_0_ maps estimated from ‘topup’ (Andersson et al., 2003) using the pre-acquired PE-reversed data. The *B*_0_ maps were motion-corrected to account for the field map orientation change due to subject motion, and then applied to each volume using the ‘FUGUE’ function in FSL (Jenkinson et al., 2012; Smith et al., 2004).

For cortical surface-based analysis, cortical reconstructions were generated automatically using Freesurfer (Desikan et al., 2006; Fischl, 2012; Fischl et al., 2002) on the MPRAGE images of each subject. A total of 9 equi-volume (Waehnert et al., 2014; Waehnert et al., 2016) cortical layers were reconstructed, and applied to the EPTI and EPI data to investigate the distribution of the z-score and the percent signal change across different cortical depths.

## 3. Results

The effect of the shot-to-shot phase variations on the EPTI reconstruction and the performance of the estimation and correction methods are evaluated in Fig. 3. Large temporal *B*_0_ variations with a range of ±10 Hz were used in this evaluation to simulate the effect at 7T. The estimated *B*_0_ variation maps (Fig. 3b) show spatial distribution similar to the reference (Fig. 3a), demonstrating the effectiveness of the *B*_0_ variation estimation method using multi-channel 1D navigators. The reconstructed images without and with correction are compared and shown for the pure SE (Fig. 3c) and a selected ASE (Fig. 3d) image. The images without any added variations are also shown in the left-most columns to illustrate baseline reconstruction errors when compared to the ground truth simulated fully-sampled data. As shown in the error maps, the pure SE image is less affected by the phase variations and presents with only a small increase in RMSEs even without correction, while the ASE image shows higher errors resulted from the phase variations due to its larger *B*_0_ phase accumulation. After correction, the errors are significantly mitigated, especially for the ASE image, and similar reconstruction accuracy is observed as in the case without any added phase variations, demonstrating the effectiveness of the proposed correction approach.

**Figure 3.**
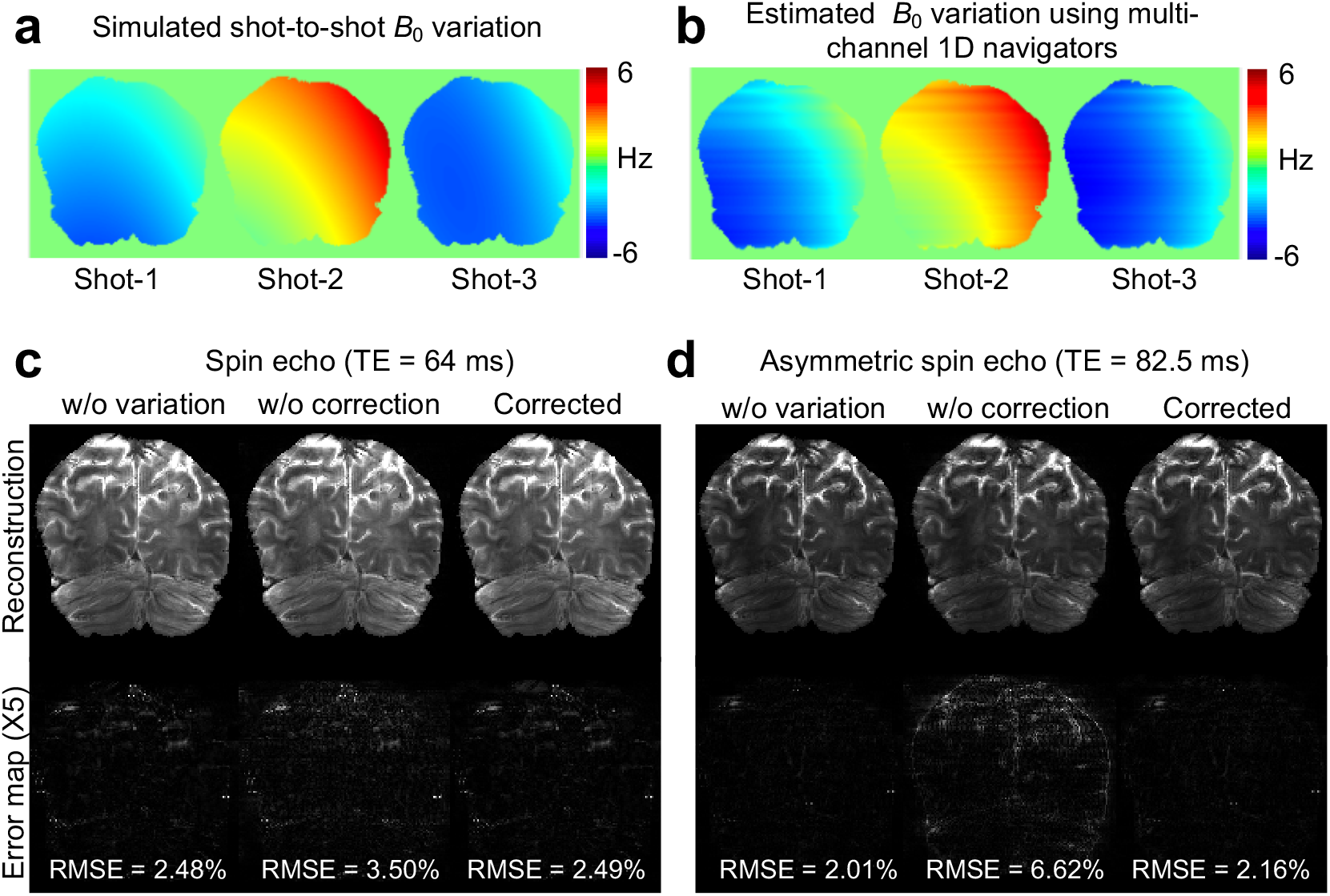
Evaluation of the shot-to-shot *B*_0_ variation estimation and correction method. (a) Simulated shot-to-shot *B*_0_ variation maps of the 3-shot acquisition (spatially 2^nd^ order). (b) Estimated *B*_0_ variation maps using multi-channel 1D navigators. (c) Reconstructed pure SE images (TE = 64 ms) and their corresponding error maps (×5) without variations, with variations but without correction, and after correction. (d) Reconstructed images and error maps for the ASE at TE = 82.5 ms. The RMSEs were listed at the bottom of each error map.

Figure 4 shows an example of the multi-echo images from a representative temporal dynamic acquired by SE-EPTI in three orthogonal views, after averaging all the runs in the visual-task experiment at 7T. The FOV of the acquisition was selected to cover the visual cortex as shown on the right. Figure 5a compares the geometric distortion between conventional SE-EPI, EPTI and EPTI-extracted EPI. Two different slices are presented with overlaid brain contours (red lines) extracted from the distortion-free TSE image collected in the same scan session. The conventional EPI shows severe distortion at multiple areas highlighted by the yellow arrows. In contrast, both EPTI and the EPTI-extracted EPI are free from such distortion artifacts and provide identical contours as the TSE reference. Note that the EPTI-extracted EPI images were generated after removing the *B*_0_ phase in the full *k*-*t* data (not possible with conventional EPI data), therefore are free from the geometric distortion, similar to the EPTI images. In addition, EPTI is also robust to dynamic *B*_0_/susceptibility changes, and provides images free from distortion changes across time as demonstrated in Fig. 5b. As shown in the zoom-in 1D signal profile along the PE direction (extracted from the locations indicated by the yellow dotted lines) across different dynamics and runs, conventional EPI suffers from dynamic changes in distortion that are hard to correct for, while the signal profiles of EPTI and EPTI-extracted EPI are almost static and consistent across time.

**Figure 4.**
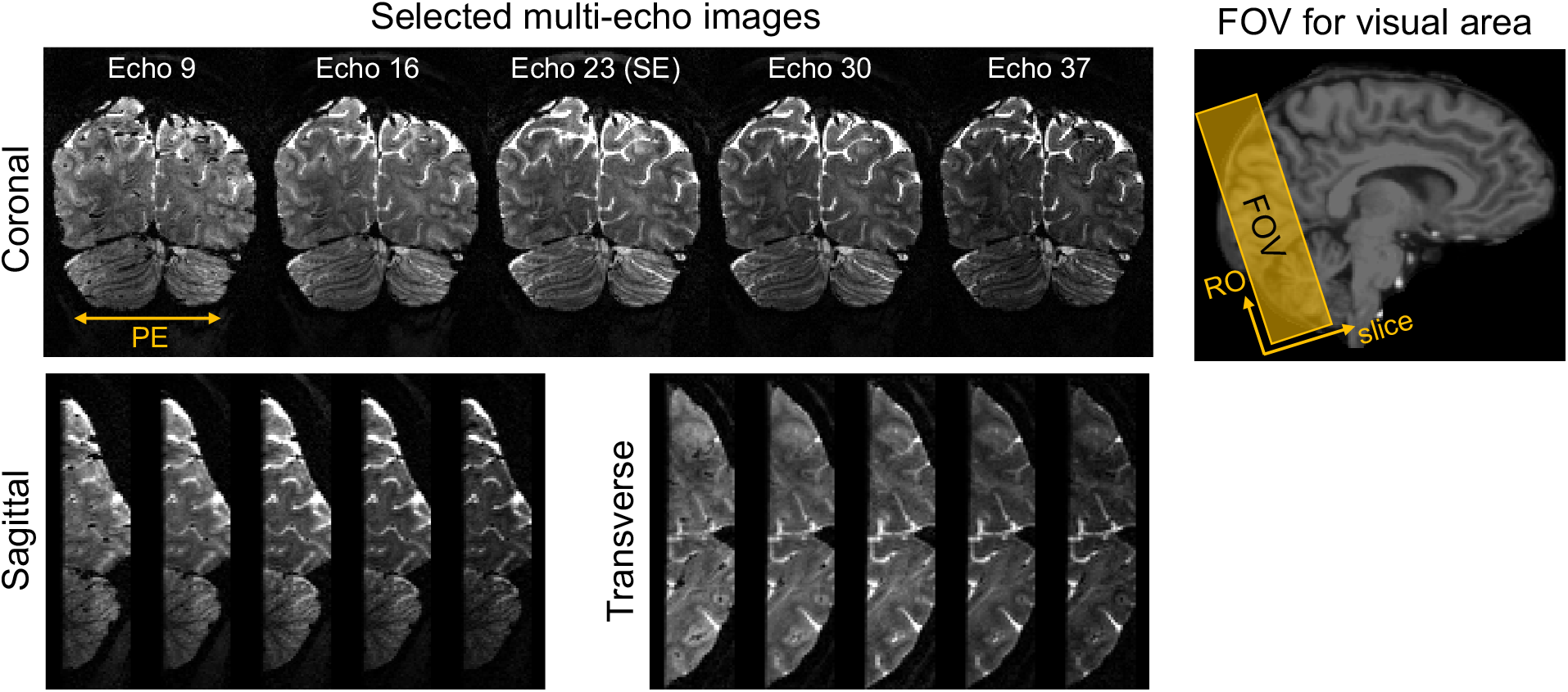
Examples of multi-echo EPTI images (left) acquired in the fMRI experiment covering the visual cortex (right). The images are shown for each dynamic after averaging all the runs. Three orthogonal views are presented for 5 selected echoes out of the total 45 echoes.

**Figure 5.**
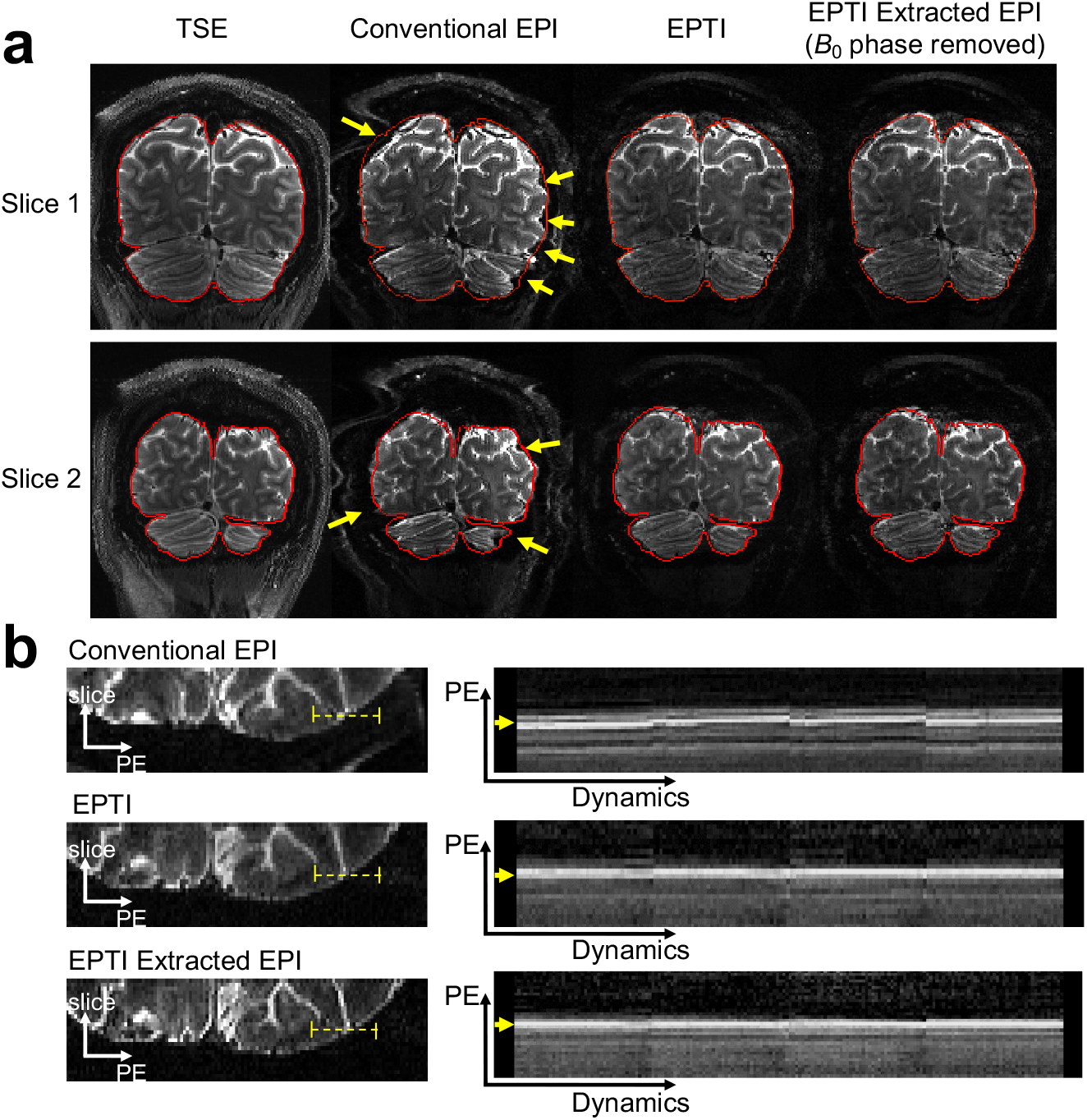
(a) Distortion comparison between the TSE reference, conventional EPI, EPTI and EPTI-extracted EPI. The image contours extracted from the TSE image are applied to all images (red lines). Conventional EPI shows obvious distortions at multiple areas (yellow arrows), while EPTI and EPTI-extracted EPI have almost identical image contours with the TSE image. (b) Evaluation of dynamic distortion changes. The zoomed-in 1D PE profiles (extracted from the locations indicated by the yellow dotted lines on the left) across different dynamics and runs are shown on the right to compare the level of dynamic distortion in conventional EPI, EPTI and EPTI-extracted EPI.

The varying T_2_′ effect across the spin-echo readout was investigated using the time-resolved multi-echo EPTI images as shown in Fig. 6. Fig. 6a shows the activation maps within visual cortex across different echoes in normalized z-scores, where a scaling was performed across different echoes based on the sum of the positive z-scores of each echo to normalize their sensitivity difference and to better compare the activation patterns. As expected, with less T_2_′ *con*trast in the images closer to the pure spin-echo (from echo 7 to echo 23, or from echo 39 backwards to echo 23), there is less activation in the CSF region (white arrows) compared to the activation in the gray matter, with the peak of the activation gradually shifting from centering in the CSF region towards the gray matter parenchyma on the side. This can also be seen in the cortical depth dependence analysis shown in Fig. 6b, where the cortical profiles of the z-score and the percent signal change across V1 of all the echoes (color-coded by echo indices, pure SE shown in green) are plotted. The profiles from different echoes were scaled to have the same value at 0.5 cortical depth as the SE (echo 23) to better compare the difference in slope. The first few echoes in blue with large amount of T_2_′ weightings exhibit the expected bias to large veins, manifesting as depth profiles that peak at the pial surface. As TE increases and moves closer to the SE from blue to green, lower and lower pial surface bias were observed in flatter depth profiles with lower values at the pial surface. At the SE position (pure T_2_), the smallest slope with minimal bias is observed. Then, as TE moves away from the SE position with more T_2_′ weighting, the pial vessel bias returns and the slope increases. In summary, the slope of these cortical profiles or the amount of large vessel bias across echoes show good correspondence to the amount of T_2_′ weighting in the theoretical signal model (Eq. 1), both of which are lowest at SE, and increase with the distance away from SE. This observation suggests that, by experimentally reducing T_2_′ weighting in the image, the macrovascular effects can be effectively reduced. It also demonstrates that EPTI provides a powerful tool to resolve all these multi-contrast images across the EPTI readout, and to investigate gradual TE-dependent BOLD signal change at a TE increment as short as 1 ms using data acquired in a single scan.

**Figure 6.**
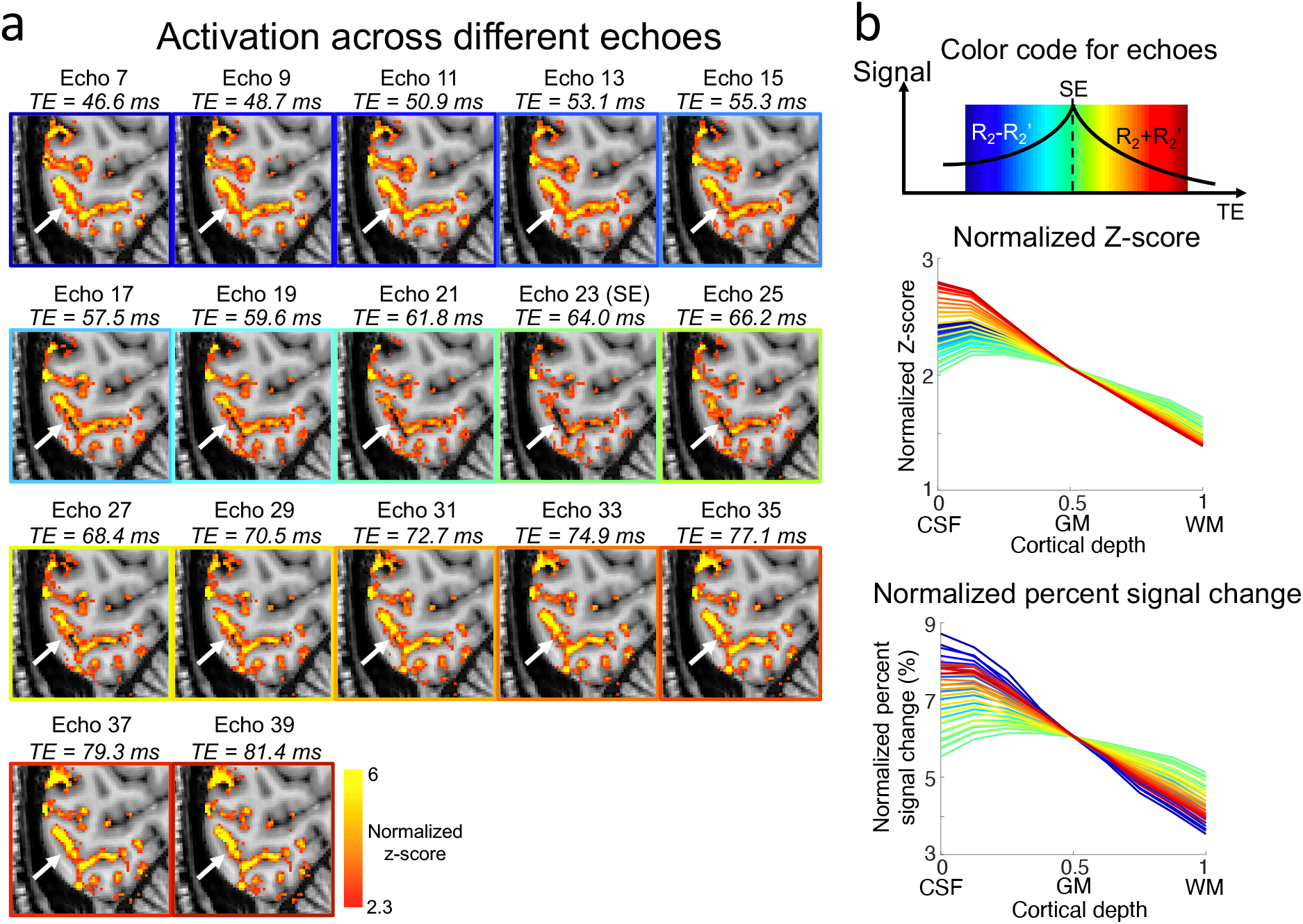
(a) Normalized z-score activation maps of different EPTI echo images. The white arrow shows an example region where the activation in the CSF is reduced relative to that in the gray matter in echoes with less T_2_′ contributions. Normalization was performed based on the sum of the positive z-scores of each echo to normalize the sensitivity differences and to better visualize activation pattern. (b) Cortical depth dependent profiles of z-score and percent signal change of all the echoes (color-coded by echo indices shown on the top) across V1. The profiles from different echoes are normalized to have the same value at 0.5 cortical depth of the SE (echo 23) to better compare the slope difference.

Figure 7 compares unnormalized activation maps (z-score) calculated from EPTI ASEs (echo 7 and 39), EPTI pure SE (echo 23), and two EPTI-extracted SE ms-EPI with ETLs of 26 ms (6-shot) and 39 ms (4-shot). The ASE images with large T_2_′ weightings show high activation in both the CSF and gray matter areas. The pure SE image and the extracted SE-EPI images show overall less activations as expected due to the reduced activation sensitivity of T_2_ contrast. Despite such a sensitivity difference, it can still be observed that the three cases of T_2_ images, including the pure SE and the two extracted SE-EPI images, show a reduction of activation in the CSF areas when compared to the gray matter’s activation level in the same maps. The pure SE image shows minimal CSF bias, pointing to EPTI’s ability to provide a more pure SE contrast with further reduction in T_2_′ contamination compared to the conventional ms-EPI, and achieve higher microvascular specificity. The cortical depth analysis of these five cases in 3 healthy volunteers are shown in Fig. 8. Both the unnormalized and normalized activity profiles are plotted. The unnormalized profiles demonstrate the sensitivity differences between T_2_′ and T_2_ weighted BOLD contrasts as previously described above, while the normalized profiles better compare the amount of large vessel bias through their slope differences. Consistent with the results in Fig. 6 and 7, the data from all the three subjects show that the pure SE image (echo 23, dark green) achieves the lowest slope and minimal macrovascular bias compared to the ASE images and the extracted conventional SE ms-EPI. In addition, the results also confirm that the conventional SE ms-EPI with a longer ETL will have larger pial vessel bias (Fig. 8, light and dark purple lines).

**Figure 7.**
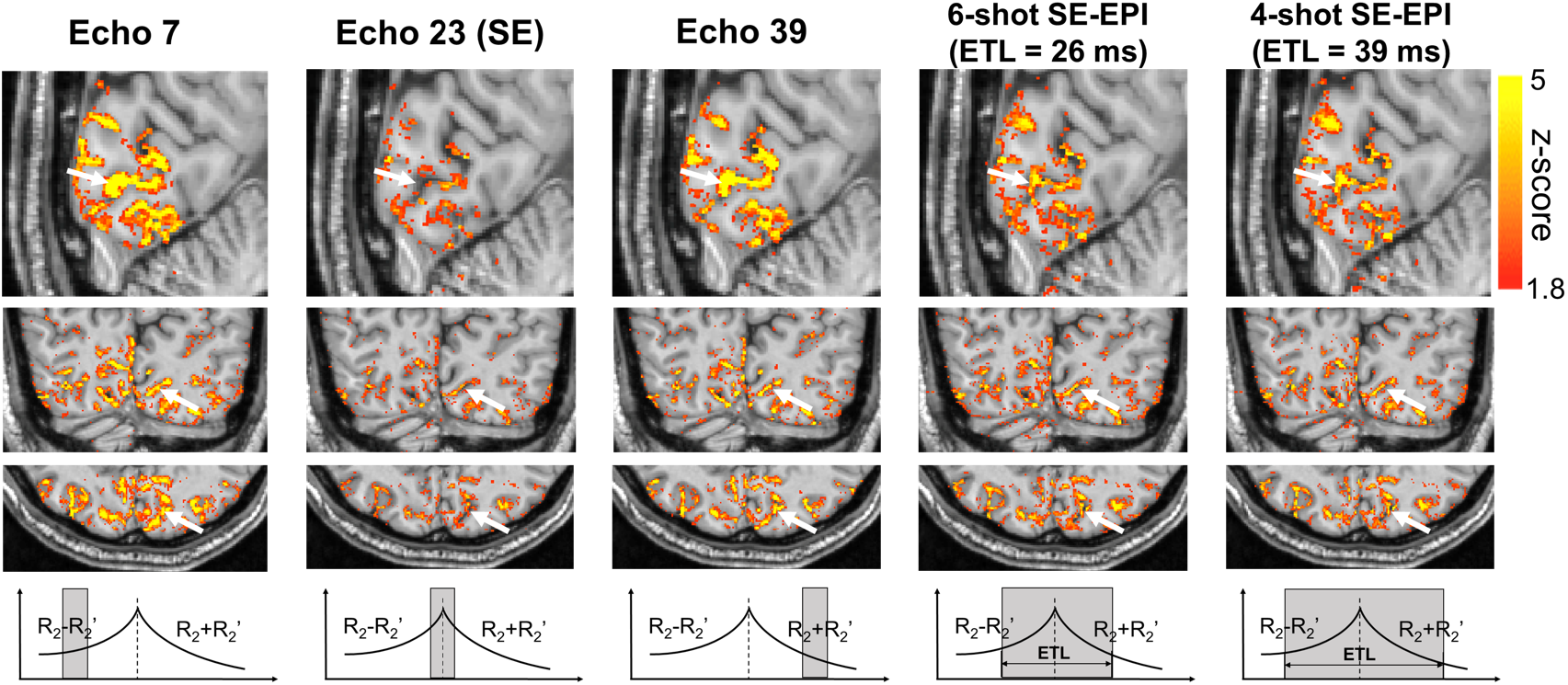
Comparison of activation in unnormalized z-score between ASE images, pure SE image, and extracted SE-EPI all obtained from EPTI. The pure SE image shows lower sensitivity than ASE echoes and the conventional SE-EPI, but despite the sensitivity difference, we can still observe that the peak of the activation map itself is more in the gray matter regions rather than centered in the CSF as indicated by the white arrows.

**Figure 8.**
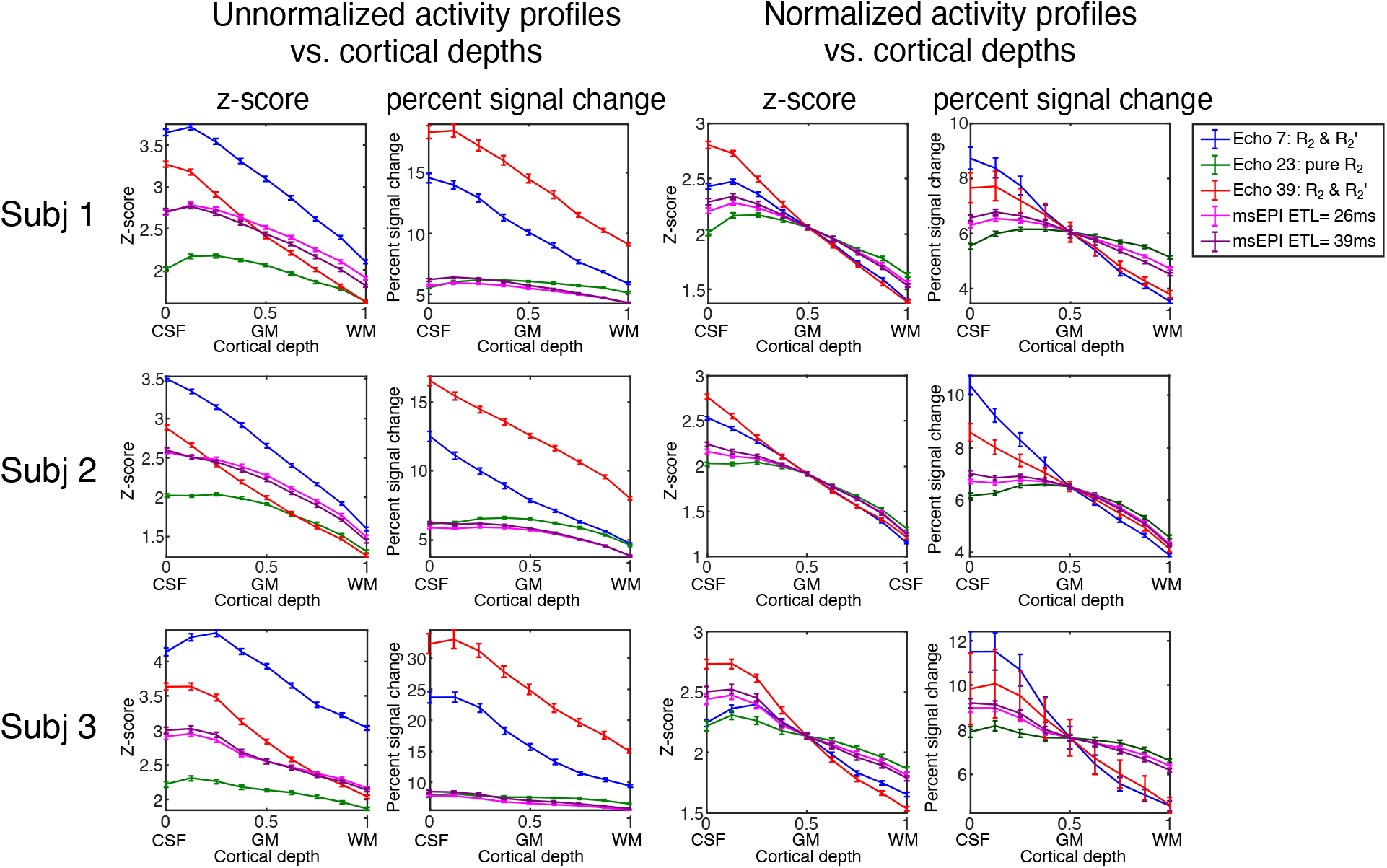
Cortical-depth profiles of the unnormalized (left two columns) and normalized (right two columns) z-score and percent signal change in three subjects (N = 3). Five EPTI data are compared in each plot, including selected ASE images (echo 7 and 39), pure SE image (echo 23), extracted SE ms-EPI with ETL = 26 ms (6 shots) and 39 ms (4 shots). In all three subjects, the pure SE image shows the lowest slope with decreased activation at the CSF-GM interface, indicating a reduced level of bias from large veins.

Further confirmation is provided by the comparison between the EPTI acquired images and the actual acquired in-plane accelerated single-shot GE and SE EPI as shown in Fig. 9. The ASE EPTI images (Echo 7 and 39) show consistent activation localization to the conventional GE-EPI, with high level of activation in both CSF and gray matter due to their large T_2_′ weighting. The cortical profile of the conventional GE-EPI also exhibits large vessel bias near the pial surface similar to the ASE EPTI images (light blue vs. dark blue or red). The conventional SE-EPI shows a reduced sensitivity to large vessels when compared to the GE/ASE acquisitions, but still has a higher activation in the CSF and a higher slope (Fig. 9a bottom row, left-most column and Fig. 9b light green) when compared to the pure SE EPTI image (Fig. 9a bottom row, right-most column and Fig. 9b dark green). This result further demonstrates the reduced draining vein effects in the pure SE EPTI image over conventional SE-EPI image. The conventional SE ss-EPI (with a GRAPPA factor of 3) also shows similar activation to the extracted SE ms-EPI with similar macrovascular bias. Noted that the conventional images were acquired in a different session than EPTI data and could have potential residual distortion after correction, which might introduce bias when comparing them with EPTI data.

**Figure 9.**
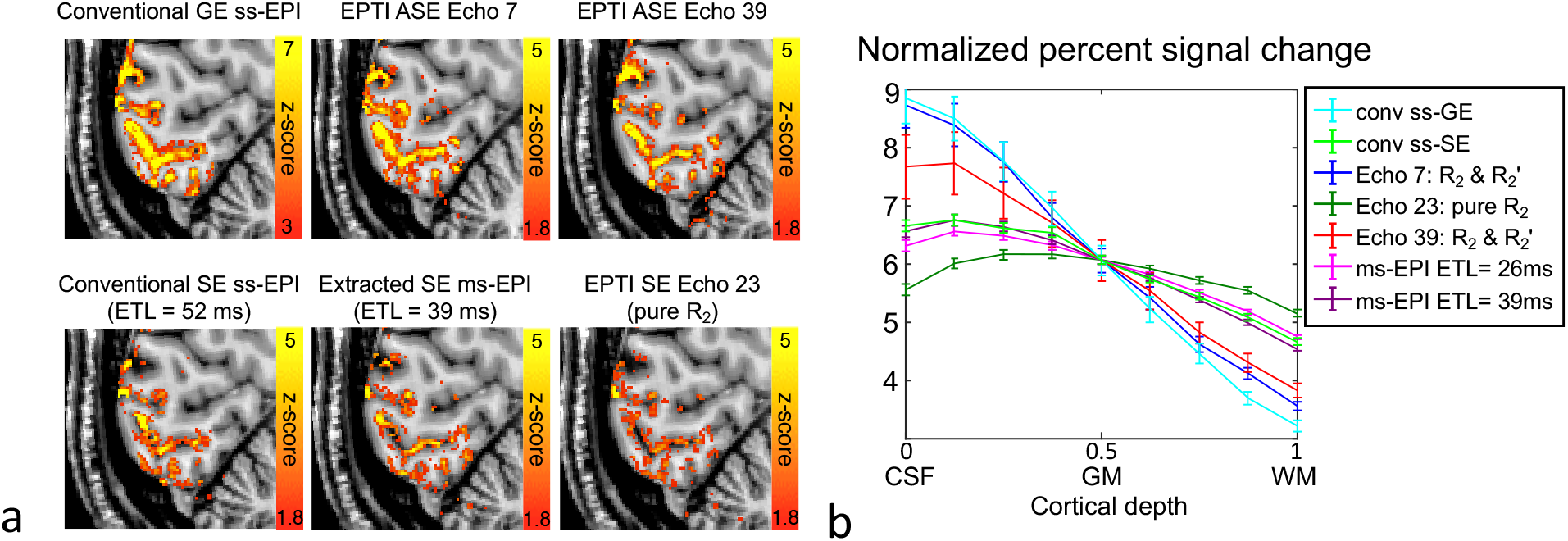
(a) Comparison between activation maps of the conventional GE-EPI, SE-EPI, and EPTI images. The ASE EPTI images (echo 7 and 39) with T_2_′ weighting show similar activation localization as the conventional GE-EPI, while the pure SE EPTI image (echo 23) provides less bias in CSF than the conventional SE-EPI. (b) Comparison of the cortical-depth profiles between the acquired GE-EPI, SE-EPI, selected EPTI echo images and EPTI-extracted EPIs with different ETLs.

## 4. Discussion and conclusions

The visual-task experiments in this study preliminarily demonstrate the ability of EPTI to time-resolve multi-echo images at a small TE increment to investigate the varying T_2_′ weighting across the spin-echo readout. The observed gradual reduction of large vein bias in image echoes closer to SE with less and less T_2_′ weighting confirms the expected association between T_2_′ contrast and the macrovascular bias. Moreover, the pure SE image acquired by EPTI shows reduced macrovascular bias compared with both the EPTI-extracted SE-EPI and the acquired conventional SE-EPI acquisition, indicating that EPTI exhibits reduced T_2_′ weighting and a purer T_2_ BOLD contrast over conventional EPI (even with multiple shots) for improved microvascular specificity. We have also experimentally confirmed that a longer ETL in SE-EPI will introduce more macrovascular bias in human experiments, consistent with the result of the previous study conducted in Monkey V1 (Goense and Logothetis, 2006). The ability of EPTI to simultaneously acquire multi-contrast images in a single scan provides a powerful tool to examine the microvascular and macrovascular contribution in T_2_ and T_2_′ BOLD contrasts. This helps to avoid any confounding differences in activation that occur between scans that would be possible in sequentially acquired runs. The similar level and consistent localization of activation seen in the EPTI ASEs data and in the GE-EPI data, and as well as between the EPTI-extracted SE-EPI data and the actual SE-EPI data (Fig. 9), demonstrates the reliability of these multi-contrast images generated from a single EPTI dataset.

In this study, we focus on investigating the reduced macrovascular contribution and improved microvascular specificity using the pure T_2_ BOLD provided by EPTI. The multi-echo images with varying T_2_′ and T_2_ BOLD contrasts, which provide varying levels of both macrovascular-sensitivity and microvascular-specificity, could also be combined to potentially allow joint modeling to improve both the sensitivity and specificity to detect neuronal activation using BOLD contrast. The multi-echo images can also be used to enhance the CNR of BOLD (Poser et al., 2006; Posse et al., 1999) or to remove physiological noise through multi-echo denoising algorithms (Kundu et al., 2012; Kundu et al., 2017; Posse et al., 1999). Moreover, while our study uses a relative long TE_SE_ (64 ms) for the EPTI acquisition to allow for extraction of the conventional EPI data with different ETLs from the same dataset for comparison, the unique time-resolved imaging approach grants the flexibility to shift the echo train to achieve shorter TEs (Wang et al., 2021b), which could be particularly useful for investigating TE-dependent signal contributions in SE-BOLD, such as varying levels of extravascular and intravascular signal changes.

Another advantage of EPTI we presented in this study is that it eliminates the geometric distortion that is common in conventional EPI-based methods due to field inhomogeneity (Fig. 5a), which is more severe at ultra-high field and in high-resolution scans. Although post-processing methods have been developed and widely used to correct the distortion, using a field map or a pair of PE-reversed acquisition, the correction for dynamic changes in distortions due to susceptibility changes resulted from multiple sources such as head motion, system drift, and respiration still remains challenging, resulting in voxels displacements between dynamics or runs and affecting the reliability of the fMRI results. In the analysis, we have performed distortion correction of conventional EPI data using field maps calculated from pairs of PE-reversed images acquired before fMRI data acquisition, and we have also accounted for the orientation change of the field map due to subject movements. However, remaining distortion changes along the PE direction are still present across different dynamics in the time-series (Fig. 5b), resulting from susceptibility and field map changes within the scan that are hard to correct using pre-scanned field maps. This can degrade the subsequent fMRI analysis. EPTI removes the geometric distortion and dynamic distortion changes through its time-resolving approach, where each echo image is formed using the data acquired at the exact same echo time (same *B*_0_ phase). The distortion-free images should help with registration to the anatomical data and lead to smaller errors in the cortical depth assignment. In addition, the time-resolving approach can also eliminate the image blurring along PE direction due to signal decay, another major limitation of conventional EPI.

Instead of geometric distortion in conventional EPI, the *B*_0_ change might lead to local blurring and elevated reconstruction errors in EPTI data as reported in our previous work (Wang et al., 2019). Here, we first characterized the impact of shot-to-shot phase variations or *B*_0_ changes on the multi-echo EPTI images. As expected, it was observed that the ASE images with more *B*_0_ phase accumulation are more sensitive to inter-shot variations than the pure SE image. To correct for this, the proposed reconstruction framework utilizes a navigator-based shot-to-shot *B*_0_ variation estimation that provides effective correction and significantly mitigates the potential blurring. A pre-reconstruction process was also incorporated to correct for higher-order *B*_0_ changes across dynamics that can further improve the reconstruction robustness.

By using the pure SE EPTI image, the cortical depth profiles of z-score and percent signal change show minimal pial surface bias. However, small amount of activation or bias was still observed within CSF or around pial surface. First, due to the use of the temporal correlation across echoes to reconstruct the highly-undersampled *k-t* data, we do not rule out the possibility that there might be some T_2_′ contrast leakage and therefore residual pial surface bias in the pure SE EPTI image. Our original work on EPTI used a GRAPPA-like reconstruction in the *k-t* space to reconstruct the images. The GRAPPA kernel interpolation along *t* could cause local temporal smoothing with an effective ETL estimated at ∼7 ms. This could cause small amount of T_2_′ contamination in the pure SE image, but is still within the range of ETLs that would provide microvasculature-dominated BOLD (Goense and Logothetis, 2006). In this work, a subspace reconstruction was employed to take advantages of the signal model prior that can accurately represent the signal evolution (errors < 0.2%), without interpolating along *t* as in the GRAPPA reconstruction, which should result in reduced amount of T_2_′ leakage and provide purer SE. To systematically characterize the potential residual T_2_′ contamination/leakage in the subspace reconstruction, a generalized approach to analyze the signal response of non-linear reconstruction (e.g., LLR-constraint subspace reconstruction in this work) might be required in the future. Second, partial volume effects and potential residual intravascular signal contribution (longer intravascular T_2_ than expected) could also be possible explanations. Biophysical modeling and simulation as well as experiments with different SE TEs (Berman et al., 2021; Pfaffenrot et al., 2021; Pflugfelder et al., 2011) should be helpful to better understand the signal contributions of SE BOLD fMRI.

At the current resolution, slightly higher activation can already be observed around the middle depth in the visual cortex, which might reflect higher microvascular density. However, the profiles are still relatively flat across depths that could possibly be explained by an averaging effect over the large V1 ROI and/or by partial volume effects (De Martino et al., 2013; Koopmans et al., 2011). Further increases in spatial resolution can help reduce partial volume effect and should provide better visualization of such laminar fMRI responses. Recent laminar fMRI studies have also shown different responses in different layers in the motor cortex (Huber et al., 2017a), which may be useful to further evaluate the EPTI approach. The temporal resolution of EPTI acquisition could also be improved to increase the statistical power of fMRI analysis. These can be achieved by further developments of our EPTI-fMRI method to enable faster *k-t* encoding, such as by incorporating low-rank modelling across the fMRI time-series (Chiew et al., 2015), utilizing more advanced *k-t* sampling trajectories (Dong et al., 2021b; Fair et al., 2020) or combining with multi-slice or multi-slab acquisition (Han et al., 2020; Setsompop et al., 2018; Setsompop et al., 2012), which will then open up an exciting opportunity to perform pure SE-fMRI at higher spatiotemporal resolution and larger spatial coverage.

## Acknowledgment

This work was supported by the NIH NIBIB (R01-EB020613, R01-EB019437, R01-EB016695, P41-EB030006, and U01-EB025162) and NIHM (R01-MH116173) and made possible by resources provided by Shared Instrumentation Grants S10-OD023637, S10-OD02363701, S10-RR023401, S10-RR023043, and S10-RR019307. We thank Azma Mareyam for providing the 64-channel phased-array coil used in this study.

